# Neural Dynamics Underlying Repeated Learning of Visual Image Sequences

**DOI:** 10.1101/2025.11.04.686589

**Authors:** Chong Zhao, Audrey Kim, Leyla Campos, Edward K. Vogel

## Abstract

Humans possess a remarkable ability to recognize visual objects with high fidelity, supported by complex neural mechanisms underlying memory retrieval. Event-related potential (ERP) studies have identified two key neural signatures of recognition memory: the parietal old/new effect and the frontal old/new effect. Despite extensive research on these ERP components, the extent to which these components reflect distinct memory processes remains debated. In the present study, we investigated how repetitive learning modulates these ERP components. Participants repeatedly studied a fixed list of 32 real-world images across up to five study-test repetitions while EEG was recorded. Additionally, a separate set size 1 condition served as a proxy for working memory. Our results showed that with increased repetitions, the parietal old/new effect exhibited enhanced amplitude and earlier peak latency, reflecting more efficient retrieval of well-learned memories. In contrast, the frontal old/new effect remained unchanged in both amplitude and timing. These findings suggest that the parietal old/new effect is a sensitive neural marker of learning-related changes in long-term memory representations, while the frontal effect is less influenced by repetition. Additionally, despite similarly high accuracy between the well-practiced set size 32 condition and the set size 1 working memory condition, both parietal and frontal old/new effects peaked significantly earlier for set size 1, suggesting that access to working memory is substantially faster than even well-practiced long-term memory. Together, our results highlight the unique role of the parietal old/new effect, but not the frontal old/new effect, in repetitive learning, despite both components being important for successful recognition of learnt visual stimuli.

## Introduction

Humans possess a remarkable capacity for recognition memory, storing visual objects with high fidelity and even spatio-temporal precision (Brady et al., 2008; Lionel, 1973; Wolfe et al., 2023). This ability prompts important questions about the cognitive and neural mechanisms that support such detailed memory. To investigate how the brain distinguishes previously encountered objects from novel ones, researchers have frequently turned to event-related potentials (ERPs) to examine the temporal dynamics of memory retrieval. ERP studies have offered valuable insights into the neural correlates of recognition memory, particularly by focusing on the differential brain responses elicited by old versus new items, commonly referred to as “old/new effects” (Kwon et al., 2023; Rugg & Curran, 2007).

The first identified old/new ERP effect is the parietal old/new effect, typically observed between 500–800 ms after stimulus onset and maximal over left parietal scalp regions. This effect is strongly associated with recollection, as it reflects greater positivity for items that are recognized with contextual detail or source memory. It is most often elicited in “remember” responses—those accompanied by high confidence and retrieval of specific episodic details—rather than in “know” responses, which reflect familiarity-based recognition with lower confidence (Friedman & Johnson Jr, 2000; Wilding & Rugg, 1996). The parietal old/new effect is consistently enhanced when participants are required to retrieve contextual features such as the source or spatial location of previously studied items (Mecklinger, 2000; Vilberg & Rugg, 2008). Neuroimaging studies have implicated the left lateral parietal cortex, particularly the angular gyrus, in the generation of this effect, suggesting its involvement in the integration and conscious experience of source information (Cabeza et al., 2008; Wagner et al., 2005). Together, these findings provide strong evidence that the parietal old/new effect serves as a neural marker of memory retrieval during recognition memory tests.

Complementing the parietal old/new effect, researchers have also identified a frontal old/new effect, which emerges earlier, typically between 300-500 ms post-stimulus, and is most prominent at mid-frontal scalp sites, and is widely associated with successful recognition. This ERP component is characterized by a more positive-going waveform for correctly recognized old items compared to new ones, particularly in conditions that emphasize familiarity. It is reliably elicited in remember/know paradigms when participants classify items as “known” rather than “remembered”, indicating a familiarity-driven process (Curran, 2000, 2004; Rugg & Curran, 2007). The amplitude of this effect is modulated by factors that influence the strength of familiarity signals, such as conceptual fluency, and priming strength (Leynes, 2012; Paller et al., 2007). Neuroimaging evidence points to regions of the anterior prefrontal cortex as potential sources of the frontal old/new effect, implicating this area in strategic monitoring and decision-making based on familiarity (Duarte et al., 2006; Yonelinas et al., 2005). Collectively, these findings support the interpretation of the frontal old/new effect as a neural correlate of successful memory retrieval during the test phase of recognition tasks.

Despite extensive research on the frontal and parietal old/new ERP effects, there remain ongoing debates about whether these two components reflect distinct neural processes or overlapping memory retrieval mechanisms. Several lines of research suggest that the two effects reflect distinct neural processes, as they respond differently to experimental manipulations and show unique temporal and topographical profiles (Curran & Cleary, 2003; Woodruff et al., 2006). For example, the frontal effect has been shown to remain intact in individuals with hippocampal damage, whereas the parietal effect appears more sensitive to tasks involving relational encoding or source retrieval (Addante et al., 2012; Opitz & Cornell, 2006). Meta-analytic findings further support this distinction, highlighting reliable differences in the timing and scalp distribution of the two effects across studies (Kwon et al., 2023). However, other work suggests that these ERP components may reflect graded or interactive processes, with overlapping contributions to recognition, particularly under conditions that vary in memory strength or decision confidence (Squire et al., 2007; Wixted, 2007). Moreover, studies combining ERP with source localization and multivariate analysis have shown both distinct and partially overlapping neural generators, indicating that the two effects may not be entirely independent (Hoppstädter et al., 2015; Rugg & Vilberg, 2013). Overall, prior studies support a functional separation between the frontal and parietal old/new effects. However, all of these studies have examined episodic memory following only a single exposure. As a result, it remains unclear how repeated learning modulates these neural signatures of successful memory retrieval.

In the present study, we addressed this question using a recognition memory learning paradigm. Our primary aim was to determine which neural signatures, the frontal or parietal old/new effects, are modulated by learning. Participants repeatedly studied a fixed set of 32 real-world images across up to five study-test cycles. This design allowed us to track how electrophysiological markers of memory change as representations become more robust through repeated exposure. We also included a portion of trials with only 1 item in the list. Because the memory load for these trials is low and the interval between study and test is uninterrupted and short, subjects likely use working memory to aid the recognition process. Considering this, we expect that the ERP old/new effects (in both frontal and parietal electrodes) from 1-item lists would be earlier than that from 32-item lists because the memorandum is already active in working memory when the test is presented and does not require the additional time-consuming process of retrieval from long-term memory. In the current study, we sought to use these trials as a comparison point for evaluating the learning effects for the 32-item lists. This would allow us to test whether repetitive learning of a 32-item list produced ERP old/new effects that begin to approximate those observed when they are supported by working memory.

## Method

### Participants

The experimental procedures were approved by the Institutional Review Board at The University of Chicago. All participants provided informed consent and were compensated with a cash payment of $20 per hour. They reported having normal color vision and normal or corrected-to-normal visual acuity. Participants were recruited from The University of Chicago and the surrounding community.

A priori power analysis was conducted to determine the required sample size to detect a repetition effect of 0.5 µV with a standard deviation of 0.5 µV, using a two-tailed paired-sample t-test. With an alpha level (α) of 0.05 and desired power (1 − β) of 0.90, the analysis indicated that a minimum of 15 subjects is required to detect the expected effect. A total of 25 subjects were recruited, and 1 subject was rejected due to excessive eye movement.

### Stimuli and Procedure

Images were selected from the THINGS dataset (Hebart et al., 2023). One exemplar was selected from each category so that no repeated category was used throughout each session. The recognition memory procedure followed established protocols in the literature and consisted of separate study and test phases (Fukuda & Woodman, 2015; Zhao, Fukuda, et al., 2025; Zhao & Woodman, 2020). In the set size 1 condition, each trial began with the presentation of a single image (5° × 5°) centered on the screen, accompanied by a white fixation cross. The image was displayed for 800 ms, during which participants were instructed to encode it into memory. Afterward, a self-paced blank interval followed, allowing participants to initiate the test phase at their own pace. Each test phase consisted of two recognition trials. On each test trial, an image appeared at the center of the screen for 800 ms along with a white fixation cross. To minimize motor and ocular artifacts, participants were instructed to avoid any movements while the fixation cross was present. After 800 ms, the fixation cross disappeared, and the image remained on screen until a response was made. One of the two test images matched the image from the encoding phase (old), while the other was a novel image that had not been seen before (new). The order of the old and new images was randomized across blocks. Participants were asked to indicate whether the image was “old” (previously studied) or “new” (not previously seen) using designated keyboard keys (“z” for old, “/” for new).

In the set size 32 condition, participants viewed a stream of 32 images, each presented sequentially at the center of the screen for 800 ms. A white fixation cross was superimposed on each image to help maintain central fixation and discourage eye movements. The interstimulus interval (ISI) between images was randomly jittered between 250 and 400 ms to reduce anticipatory responses. Following the encoding phase, participants completed a series of recognition test trials. On each test trial, a single image was displayed at the center of the screen for 800 ms, again with a central white fixation cross. To minimize motor and ocular artifacts, participants were instructed to remain still and refrain from any movements while the fixation cross was visible. After 800 ms, the fixation cross disappeared, and the image remained on screen until a response was made. Half of the test images had been shown during the preceding study phase (old), while the other half were entirely new (new). The order of old and new images was randomized across trials. Participants indicated whether each image was old or new using the same response keys as in the set size 1 condition (“z” for old, “/” for new).

To investigate the effect of repetition-based learning, the study and test phases of the set size 32 condition were repeated five times with pseudorandomized sequence without any constraint in shuffling old and new images. Across these five repetitions, the old images remained the same, allowing participants to learn them across repeated exposures, while new images were replaced with novel ones on each test phase to avoid familiarity confounds (see **Fig. 1**). For recognition memory, we use accuracy (%) as our primary behavioral measure.

**Fig 1.**
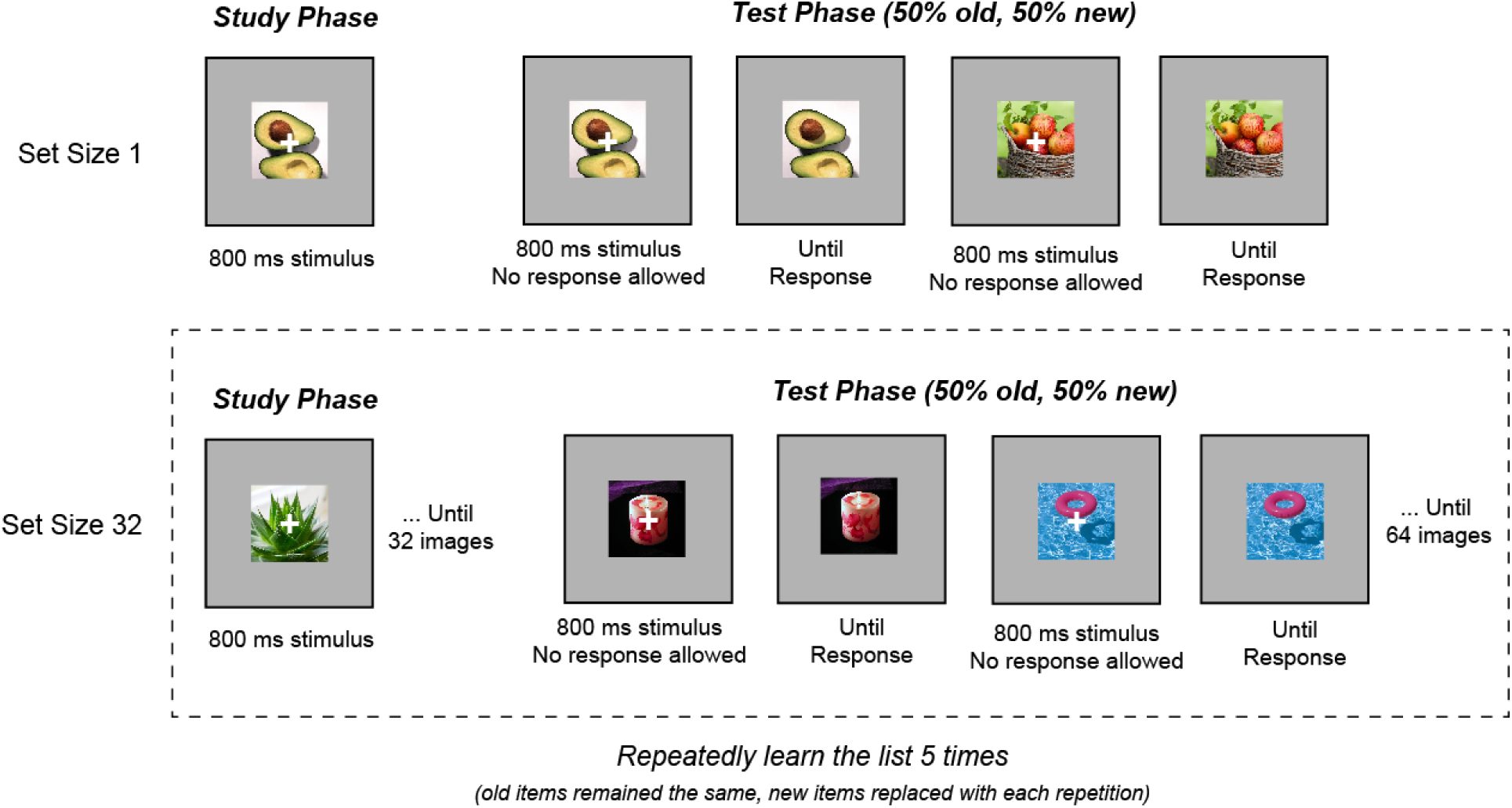
Recognition Memory and Learning Paradigm. In the *set size 1* condition, each trial began with a single centrally presented image (5° × 5°) shown for 800 ms with a white fixation cross. Participants encoded the image into memory while maintaining fixation. A self-paced blank screen followed, after which participants completed two recognition trials. Each test image appeared at the center for 800 ms with a fixation cross, followed by a response window where the image remained on screen until a response was made. One image was old (from the encoding phase), and the other was new; their order was randomized. Participants responded using keyboard keys (“z” = old, “/” = new). In the *set size 32* condition, participants viewed a stream of 32 images, each presented for 800 ms with a superimposed fixation cross. Interstimulus intervals were jittered between 250–400 ms. After the stream, participants completed a series of recognition test trials following the same timing and response procedure as in the set size 1 condition. To examine repetition-based learning, the study-test cycle was **repeated five times**. The same 32 images were used as old items across repetitions, while new images were replaced on each test phase to prevent familiarity confounds.

### Preprocessing and artifact rejection of EEG signals

Participants were positioned in an electrically isolated booth, with their heads stabilized using a cushioned chinrest placed 74 cm from the display screen. Electroencephalographic (EEG) signals were recorded via 30 active silver/silver chloride (Ag/AgCl) electrodes integrated into a stretchable cap (actiCHamp system, Brain Products, Munich, Germany), arranged according to the international 10-20 placement standard (electrode sites: Fp1, Fp2, F7, F8, F3, F4, Fz, FC5, FC6, FC1, FC2, C3, C4, Cz, CP5, CP6, CP1, CP2, P7, P8, P3, P4, Pz, PO7, PO8, PO3, PO4, O1, O2, Oz). Additional electrodes were adhered to the left and right mastoids using adhesive stickers, and a ground electrode was integrated at the Fpz site within the cap. EEG signals were initially referenced to the right mastoid and subsequently re-referenced offline to the average of both mastoids. The signal was bandpass filtered (0.01–80 Hz) with a 12 dB/octave roll-off and digitized at a sampling rate of 500 Hz. All electrode impedances were maintained below 10 kΩ.

To track ocular activity such as blinks and saccades, both electrooculographic (EOG) and eye-tracking data were recorded. EOG was captured using five passive Ag/AgCl electrodes: two for vertical EOG (above and below the right eye), two for horizontal EOG (approximately 1 cm lateral to each eye), and one ground electrode on the left cheek. Eye position was also monitored with a desktop-mounted EyeLink 1000 Plus system (SR Research, Ontario, Canada), operating at a sampling rate of 1,000 Hz.

### Artifact Rejection

For horizontal eye movements, we employed a sliding-window algorithm to identify horizontal eye movements using both horizontal electrooculogram (HEOG) signals and eye-tracking gaze data. For the HEOG-based detection, a split-half sliding-window method was applied with a 100 ms window moving in 10 ms increments. An eye movement was flagged if the voltage difference between the two halves of the window exceeded 20 µV. This HEOG-based artifact detection was used only in trials where eye-tracking data were unreliable or unavailable. In parallel, eye-tracking rejection was carried out by analyzing horizontal (x-axis) and vertical (y-axis) gaze positions using the same 100 ms window and 10 ms step size. A trial was excluded if eye position shifted more than 0.5° of visual angle within any given window.

To detect blinks, we applied a sliding-window analysis to the vertical EOG signal using an 80 ms window and a 10 ms step size. A blink was marked when the voltage change across the window exceeded 30 µV. Complementarily, we identified blink periods in the eye-tracking data by locating segments where positional data were missing, indicating that the eyes were closed.

### Frontal and Parietal Old/New Effect Analysis

To calculate the parietal and frontal old/new effects, we first identified sets of electrodes associated with each region. Based on prior literature, for the parietal old/new effect, we included electrodes PO3, PO4, PO7, PO8, P3, P4, P7, P8, and Pz. For the frontal old/new effect, we used electrodes F3, F4, F7, F8, and Fz. For each region, we computed the average event-related potential (ERP) time series across the selected electrodes to obtain a representative waveform. We then focused on correct trials within each set size and repetition condition and categorized them into two trial types: old (images that participants had previously studied) and new (novel images that had never been studied). We computed the average ERP for each trial type separately. The old/new effect was then derived by subtracting the ERP waveform for new trials from that for old trials, calculated separately for each participant. This difference waveform, named old/new effect, reflects neural activity specific to recognition memory.

## Result

Our primary goal was to examine how repetition influences the neural signatures of memory. To do so, we first needed to confirm that participants were indeed learning the images through repeated exposure. Before looking at the learning effect, we first verified the well-established list length effect using two baseline conditions, set size 1 and set size 32, with no repetitions. Participants showed significantly lower recognition accuracy when encoding 32 images compared to just 1 (*t*(23) = 6.80, *p* < .001, **Fig. 2**). Next, we examined whether participants’ accuracy improved with repeated exposure to the 32-image set, which would indicate learning across repetitions. As predicted, accuracy significantly increased over the five study-test cycles (*F*(4, 18) = 16.78, *p* < .001). Post-hoc t-tests revealed that accuracy improved substantially from the first to the second repetition (*t*(23) = 6.80, *p* < .001), with a modest but statistically marginal gain from the second to the third repetition (*t*(23) = 2.04, *p* = .05). No further significant improvements were observed between the third, fourth, and fifth repetitions (all *p*s > .30).

**Fig 2.**
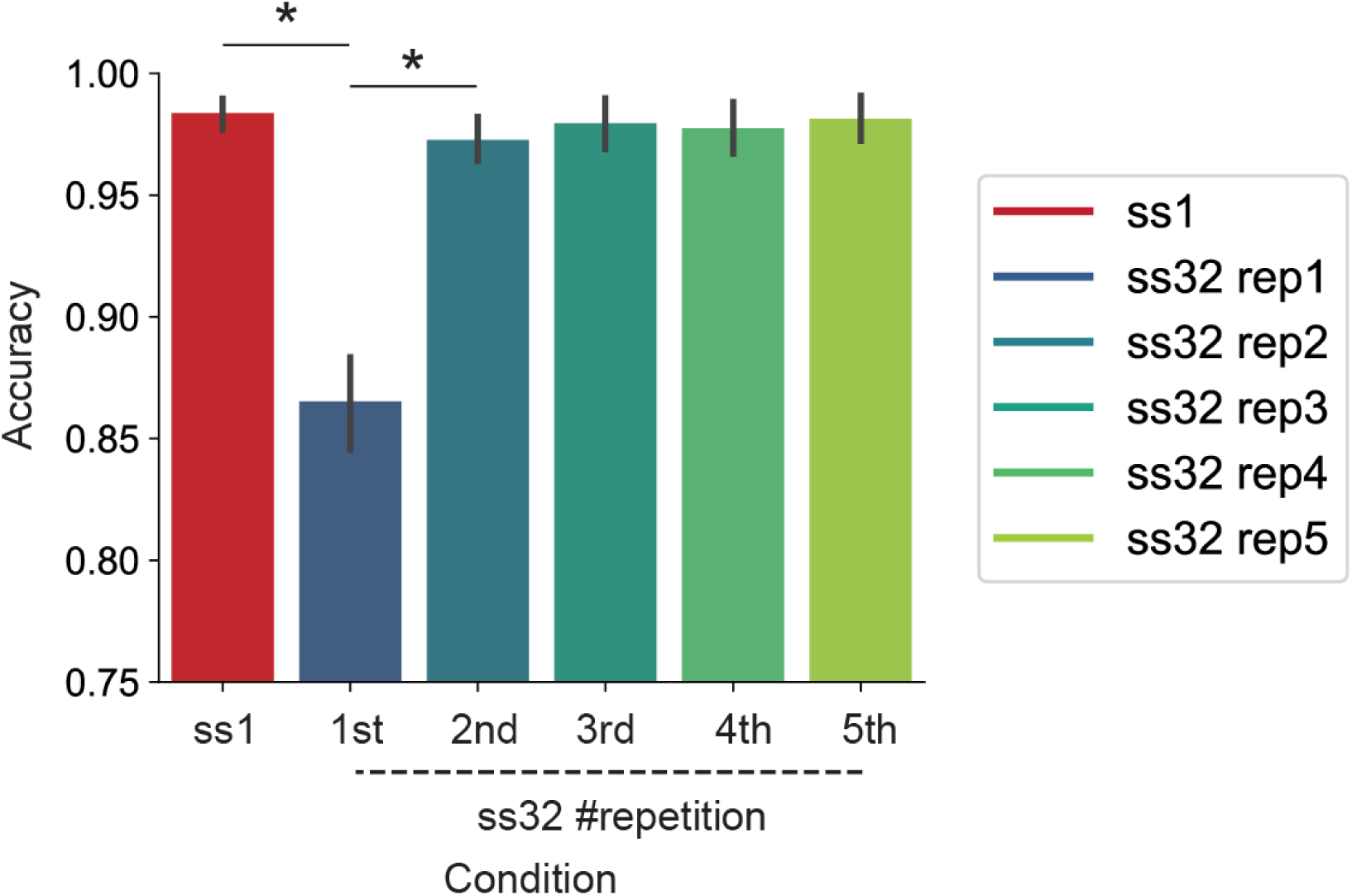
*Recognition Memory and Learning Behavioral Accuracy.* Participants showed improved accuracy from repetition 1 to 2 in the set size 32 condition, and accuracy in later repetitions did not differ significantly from that of the set size 1 condition.

Overall, the behavioral results indicate that although participants had a lower accuracy with larger memory loads (set size 32 versus set size 1), they showed clear evidence of learning, as accuracy improved with repeated exposure to the set size 32 list. After confirming that participants were indeed learning the image list, we turned to the EEG data to examine which of the neural signatures of memory were modulated by this learning. Our first focus was the parietal old/new effect, a well-established ERP component associated with recognition memory (see **Fig. 3**). According to prior research, a greater positivity in the ERP signal for correctly recognized old items (compared to new ones) reflects stronger memory (scalp topography shown in **Fig. 5**). To quantify this, we measured the mean amplitude of the parietal old/new effect within a 200–600 ms window following test stimulus onset. We found that this neural signature increased with repetition (*F*(4, 18) = 4.24, *p* = .003, BF_10_ = 24.83). Specifically, the effect grew significantly from the first to the third repetition (*t*(23) = 2.27, *p* = .03, BF_10_ = 1.81), and again from the third to the fourth (*t*(23) = 2.36, *p* = .03, BF_10_ = 2.12). However, no further increase was observed between the fourth and fifth repetitions (*t*(23) = 0.23, *p* = .82, BF_10_ = 0.22). These findings suggest that repeated study of the same image list leads to a strengthening of the parietal old/new effect. Additionally, we showed that the parietal old/new effect for set size 1 was similarly high as set size 32 repeated 4 or 5 times (set size 1 vs. set size 32 repeated 4 times: *t*(23) = –1.24, *p* = 0.23; size 1 vs. set size 32 repeated 5 times: *t*(23) = –0.88, *p* = 0.39), suggesting that repetition strengthened well-trained long-term memory recollection to a similar level as working memory.

**Fig 3.**
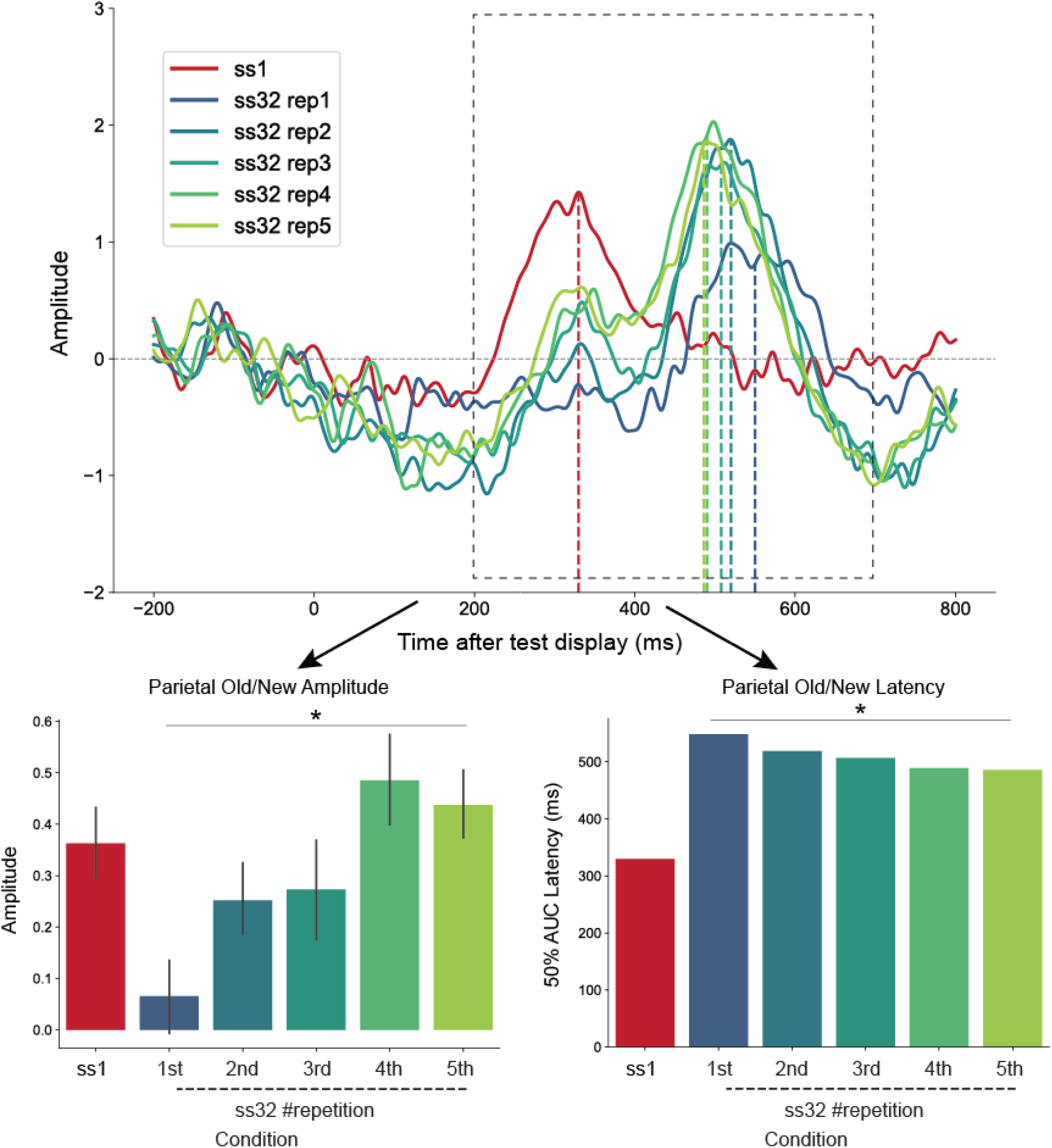
*Amplitude and Latency of Parietal Old/New Effect.* A higher amplitude and earlier latency of the parietal old/new effect were observed with increased repetitions. However, despite the latency in the set size 32 condition becoming earlier across repetitions, it remained significantly later than in the set size 1 condition.

**Fig 4.**
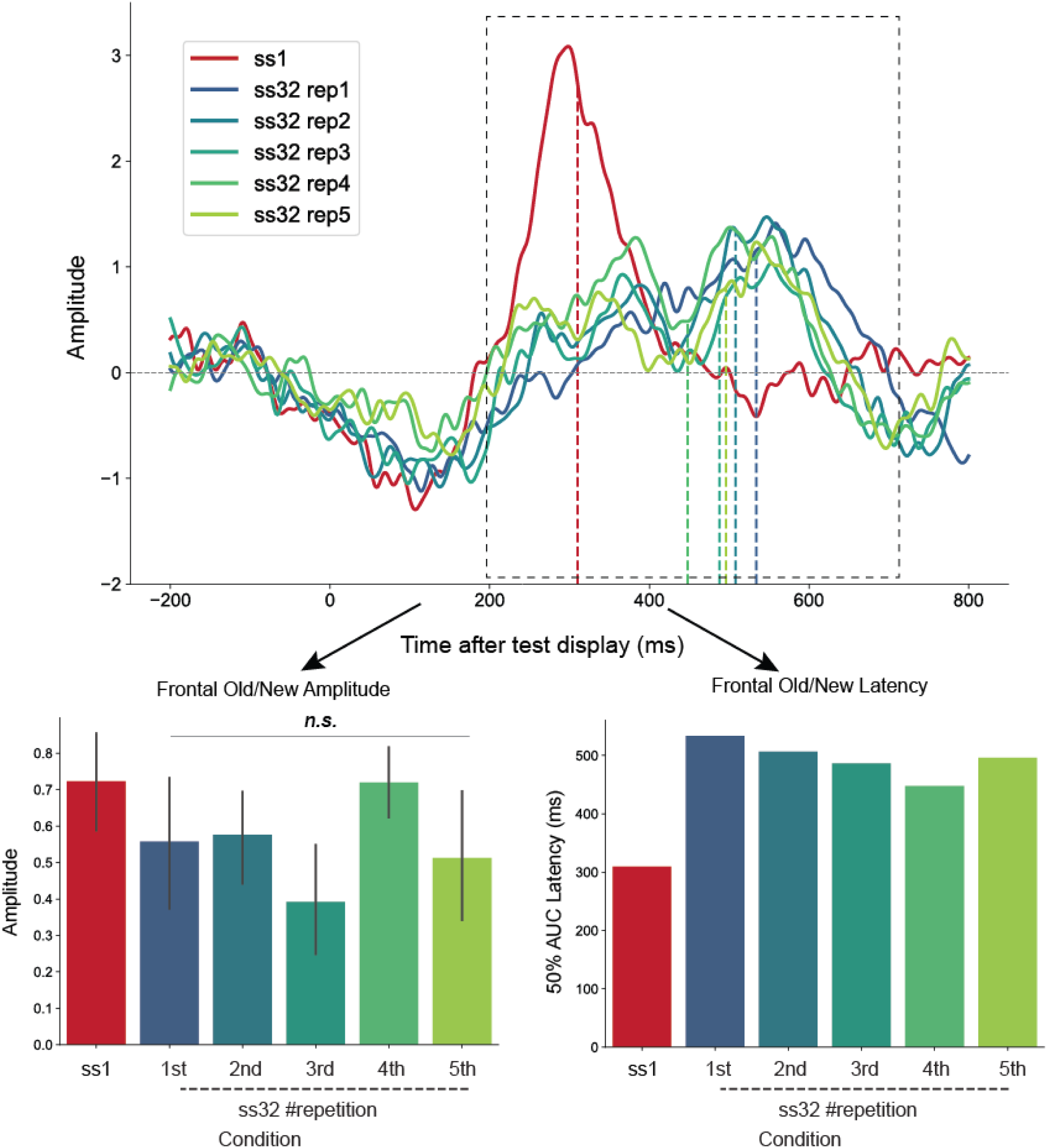
*Amplitude and Latency of Frontal Old/New Effect.* In contrast to the parietal old/new effect, the amplitude and latency of the frontal old/new effect did not change significantly across repetitions. However, despite the latency in the set size 32 condition becoming earlier across repetitions, it remained significantly later than in the set size 1 condition.

**Fig 5.**
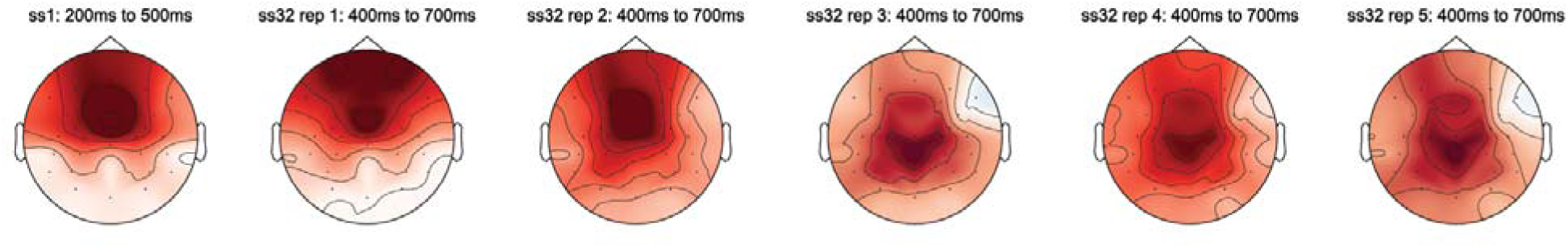
Scalp Topography of set size 1, and set size 32 repetition 1 to 5 conditions.

In addition to increasing amplitude, previous research has shown that earlier latency of the parietal old/new effect is associated with faster memory-based decision-making during the test phase. To examine this, we conducted a 50% area-under-the-curve (AUC) analysis (Kiesel et al., 2008), identifying the time point at which half of the total ERP area was reached within the relevant time window. To maximize reliability, we employed a jackknife approach for latency estimation, using a leave-one-out method across 24 participant subgroups. We calculated the latency for each subgroup and then averaged these values to obtain a robust estimate of the component’s latency. In line with our findings on amplitude, we observed that repetition significantly reduced the latency of the parietal old/new effect. Specifically, latency decreased with each successive repetition (*F*(4, 18) = 10.57, BF_10_ = 1.69 * 10^4^, *p* < 0.001): from 549.68 ms on the first exposure to 519.44 ms on the second, 506.96 ms on the third, 488.88 ms on the fourth, and 485.44 ms on the fifth (all *p*s < .001). These results indicate that repetition not only strengthens the amplitude of the parietal old/new effect but also accelerates its onset, reflecting more efficient memory retrieval with repeated exposure.

Another well-established neural marker of successful memory is the frontal old/new effect, whose amplitude has been linked to increasing memory strength, explicit memory retrieval, and even priming (scalp topography shown in **Fig. 5**). Because prior research suggests that this component becomes more positive with stronger memory signals, we examined whether repeated learning would enhance the frontal old/new effect in a manner similar to the parietal old/new effect. Contrary to our expectations, we found no evidence that repetition increased the amplitude of the frontal old/new effect (*F*(4, 18) = 0.58, *p* = .68, BF_10_ = 0.036, see **Fig. 4**). Paired t-tests comparing all consecutive repetitions also showed no significant changes in amplitude across repetitions (all *p*s > .60). Additionally, the frontal old/new effect observed in the set size 1 condition did not differ significantly from any repetition of the set size 32 condition (all *p*s > .24). When replicating our finding with non-parametric tests (Sawaki, Geng, & Luck, 2012; Maris & Oostenveld, 2007; Sawaki & Luck, 2013), we again found that all set size 32 repetitions and set size 1 frontal old/new effect amplitude were mostly the same (set size 1 not different from set size 32 repetition #1,2,4 and 5; only higher than repetition #3, corrected *p* = 0.03), despite its high peak and very steep rise-to-peak compared to set size 32 conditions. These findings suggest that, unlike the parietal old/new effect, the frontal old/new effect is not modulated by repeated exposure to the same stimuli, indicating that repetition does not strengthen this particular neural signature of memory.

In addition to examining the amplitude of the frontal component, we also analyzed the latency of the frontal old/new effect during the test phase. Using the same method applied to the parietal old/new latencies, we conducted a 50% area-under-the-curve (AUC) analysis to identify the time point at which half of the total ERP area was reached within the relevant time window. To ensure robust latency estimates, we employed a jackknife approach with a leave-one-out method across 24 participant subgroups, calculating and then averaging the latency for each subgroup.

In contrast to the parietal old/new effect, the latency of the frontal old/new effect did not show a reliable change across repetitions (*F*(4, 18) = 0.64, BF_10_ = 0.042, *p* = .63). Although there was an initial reduction in latency, from 533.04 ms on the first exposure to 506.08 ms on the second, and 487.28 ms on the third, this trend did not continue. Instead, latency plateaued at 446.84 ms on the fourth repetition and slightly increased to 495.04 ms on the fifth. This non-monotonic pattern suggests that the frontal old/new effect is unlikely to be the primary mechanism driving the benefits of repetition during learning. Rather, the absence of a consistent latency reduction implies that other neural processes may be more directly responsible for the observed improvements in memory retrieval efficiency. Furthermore, all of these latencies were significantly slower than that observed for the frontal old/new effect at set size 1 (308.88 ms), indicating that while this component is generally sensitive to memory access time, it does not exhibit the same orderly change with repetition as the parietal old/new effect. Lastly, we asked whether recency in a working-memory list reflects a stronger recollective experience, indexed by the parietal old/new effect, or instead a familiarity boost, indexed by the frontal old/new effect. To test this, we split the set-size-1 trials into two types: (1) “old-first” trials, where the old image was presented first at test immediately after encoding, and (2) “new-first” trials, where the new image was presented first and the old image came second. We found a stronger parietal old/new effect for old-first set-size-1 trials (*t*(23) = 2.09, *p* = 0.047), but no corresponding frontal old/new effect (*t*(23) = 0.64, *p* = 0.53), relative to new-first trials. This suggests that the recency of an experience, when no interfering distractor (i.e., new image in our task) intervenes, can significantly enhance recollection, but not familiarity, for a working-memory representation.

**Fig 6.**
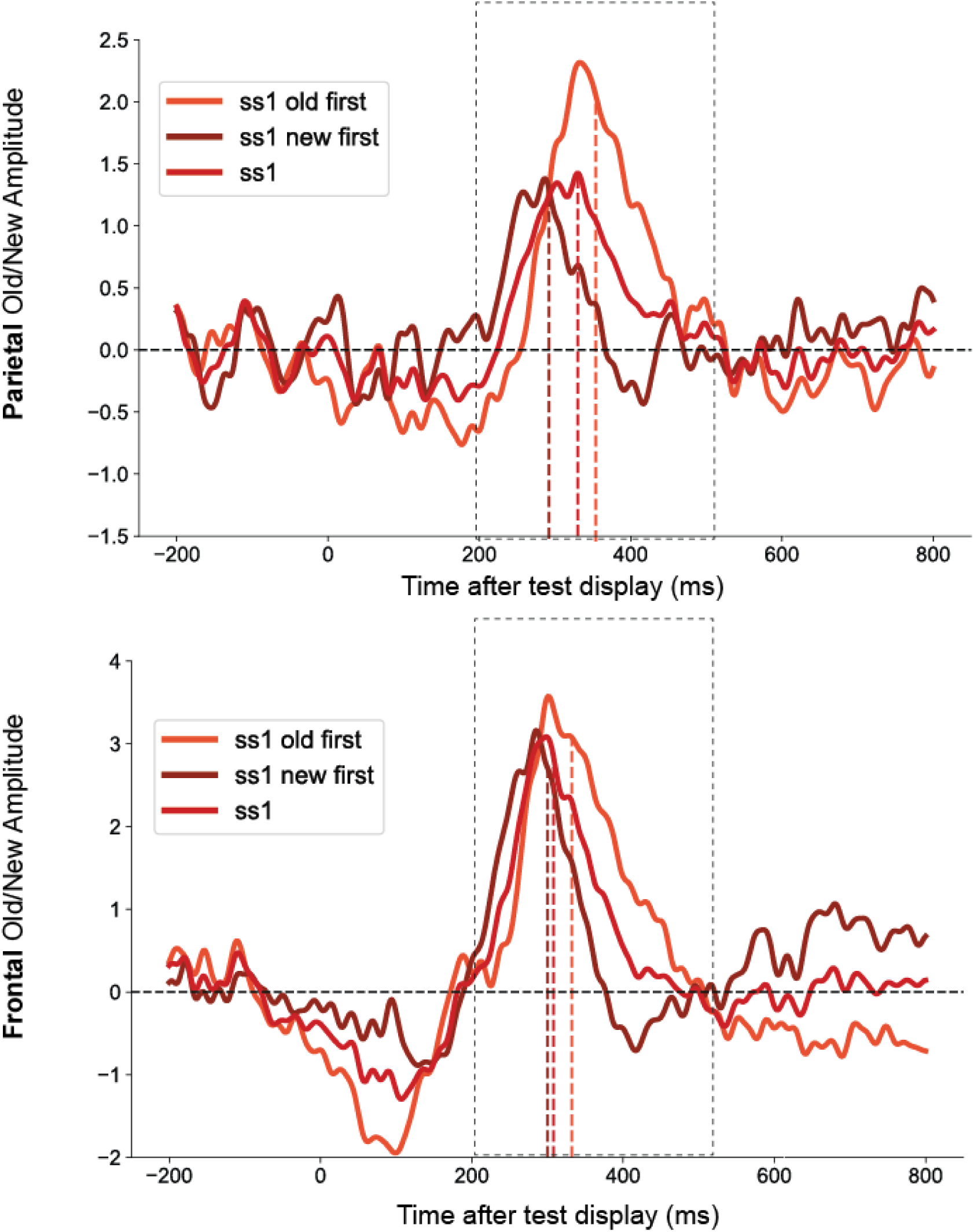
*Parietal and Frontal Old/New Effect for set size 1: serial position effect.* Within set size 1 condition, set size 1 old first trials (encoding and test had 0 image in between) triggered a particularly high parietal old/new effect amplitude compared to set size 1 new first trials, but showed no differences in frontal old/new amplitude. Q

## Discussion

In our current study, we aimed at quantifying the neural correlates of repetitive learning in the human brain. We asked the participants to learn a sequence of 32 images repetitively for up to 5 times while EEG was recorded. In particular, we are interested in whether parietal old/new effect and frontal old/new effect, two ERP correlates of recognition memory test, change with repetitive learning. We found that the parietal old/new effect increased in its amplitude and the peak became earlier with more repetitions. By contrast, we found that neither the amplitude nor the latency of frontal old/new effect was impacted by number of repetitions. These results suggest a dissociation between the two old/new effects: one, the parietal old/new effect, that becomes earlier and larger through repeated learning; and another, the frontal old/new effect, that appears largely the same despite significant behavioral learning improvements through repeated practice.

Our study demonstrated that two well-established ERP components, the parietal and frontal old/new effects, exhibit distinct patterns of change as participants repeatedly study a fixed sequence of images. Prior research has consistently linked the parietal old/new effect to recollection and the frontal old/new effect to familiarity (Kwon et al., 2023; Rugg & Curran, 2007; Yonelinas, 1994). However, the functional specificity and dynamic interaction of these processes during repeated learning remain incompletely understood. Our findings suggest that study repetition provides a powerful lens through which to dissociate these memory processes. As memory representations become more robust with repeated exposure, the changing dynamics of the frontal and parietal ERP components may reflect shifts in the relative contributions of familiarity and recollection. Importantly, the dissociation observed in our data supports the idea that these two processes follow different trajectories during sequence learning.

First, our study showed that repeating a sequence selectively increased ERP markers of recollection, but not familiarity. These neural findings align with prior behavioral evidence that, under non-speeded test conditions, item repetitions selectively increase recollection rates (Jacoby, Jones, & Dolan, 1999). Moreover, our results are consistent with empirical work showing that increasing item repetitions produces larger gains in recollection than in familiarity estimates in recognition memory tasks (Parkin & Russo, 1993; Parkin, Gardiner, & Rosser, 1995). In contrast to the parietal recollection findings, we did not observe a significant change in the frontal old/new effect with repetition. One possibility is that a single long exposure to an image is already sufficient to saturate the familiarity signal. This aligns with the picture-superiority effect observed in studies using images rather than words (Rowe & Paivio, 1971). Prior behavioral work has also shown that familiarity responses can occur even when recollective detail is not retrieved by simply accessing the picture’s name (Dewhurst & Conway, 1994). Given this relatively low threshold for familiarity with images, additional repetitions may not enhance familiarity to the same extent that they enhance recollection. Future studies should continue to leverage repeated learning designs to clarify how recollection and familiarity contribute to long-term memory formation, and how these processes are reflected in neural activity over time.

Additionally, our study addressed a key secondary question: how well-learned long-term memories differ from working memory in their neural representation. To examine this, we included a block of set size 1 image recognition trials, designed to primarily engage working memory processes, given that memory loads below three or four items typically fall within working memory capacity (Cowan, 2001; Luck & Vogel, 1997; Vogel & Machizawa, 2004). We focused on the timing (latencies) of ERP components associated with recognition. Notably, both the frontal and parietal old/new effects occurred significantly earlier for the set size 1 condition compared to the well-learned set size 32 sequence, even though participants exhibited near-ceiling accuracy for both conditions. This temporal dissociation suggests that robust episodic long-term memories, despite their strength, are neurally distinct from working memory representations. These findings align with prior work examining proactive interference and the separability of memory systems (e.g., Nee et al., 2008; Oberauer, 2009), and extend that literature by showing that ERP latencies can provide a sensitive neural marker for distinguishing between working memory and long-term memory. Future research should further explore this distinction by increasing the number of learning repetitions and extending the study period across multiple sessions (Adam et al., 2024; Musfeld et al., 2023). Such designs would allow investigation into the role of sleep-dependent consolidation and whether expertise-like memory representations begin to resemble working memory in either timing or topography. Notably, we also discovered that recency effect in working memory, where no new image happens between encoding and test, produced a particularly high parietal old/new effect, suggestive of a high level of recollection.

Lastly, we found that the latency of the parietal old/new (P3) effect decreased with repeated learning. This suggests that learning may allow participants to compress or restructure larger memory sets into more efficient representations. This finding aligns with prior research in working memory, where higher set sizes are associated with delayed P3 responses due to increased decision-making demands (Hyun et al., 2009). Extending this to long-term memory, our data showed that recognition responses for set size 32 were significantly slower than for set size 1, despite comparable accuracy, indicating that retrieving from larger sets involves more prolonged or effortful processing. Critically, the reduction in P3 latency with practice highlights this component as a potential marker of retrieval efficiency. Similar to findings in working memory, where P3 latency reflects the efficient editing of relevant items (Zhao, Adekoya, et al., 2025), our results suggest that learning may involve a comparable editing process in long-term memory. Future studies should further investigate how such editing mechanisms operate across varying memory loads and timescales, and whether additional practice enables more efficient compression of larger memory sets into smaller, more manageable representations. Moreover, well-established long-term memory representations may also alter the speed of motor planning. This can be indexed by known ERP components such as the lateralized readiness potentials (Zhao, Adekoya, et al., 2025) during recognition. Future work should therefore investigate how motor planning changes as expertise develops (Ericsson et al., 1993; Ericsson & Kintsch, 1995; Zhao & Vogel, 2025), as well as how learning shapes motor working memory.

## Notes

### Competing Interest Statement

The authors have declared no competing interest.

### Funding

- National Institute of Mental Health: ROIMH087214
- Office of Naval Research: N00014-12-1-0972, N00014-23-1-2768

